# Scaling of stochastic growth and division dynamics: A comparative study of individual rod-shaped cells in the Mother Machine and SChemostat platforms

**DOI:** 10.1101/2023.11.23.568485

**Authors:** Karl F Ziegler, Kunaal Joshi, Charles S Wright, Shaswata Roy, Will Caruso, Rudro R Biswas, Srividya Iyer-Biswas

## Abstract

Microfluidic platforms enable long-term quantification of stochastic behaviors of individual bacterial cells under precisely controlled growth conditions. Yet, quantitative comparisons of physiological parameters and cell behaviors of different microorganisms in different experimental and device modalities is not readily possible owing to experiment-specific details affecting cell physiology in confounding ways. To rigorously assess the effects of mechanical confinement, we designed, engineered, and performed side-by-side experiments under otherwise identical conditions in the Mother Machine (with confinement) and the SChemostat (without confinement), using the latter as the ideal comparator. We established a protocol to cultivate a suitably engineered rod-shaped mutant of *Caulobacter crescentus* in the Mother Machine, and benchmarked the differences in stochastic growth and division dynamics in the Mother Machine with respect to the SChemostat. While the single-cell growth rate distributions are remarkably similar, the mechanically confined cells in the Mother Machine experience a substantial increase in interdivision times. However, we find that the division ratio distribution precisely compensates for this increase in the interdivision times, which in turn reflects identical emergent simplicities governing stochastic intergenerational homeostasis of cell sizes across device and experimental configurations, provided the cell sizes are appropriately mean-rescaled in each condition. Our results provide insights into the nature of the robustness of the bacterial growth and division machinery.

## Introduction

Time-lapse microscopy is an essential technique for the study of dynamic and stochastic biological processes in individual bacterial cells. One of the key challenges in conducting such experiments lies in the creation of an environment that is suitable for the microculture of bacteria (including replenishment of nutrients) while also permitting high-resolution microscopy (including compatibility with common imaging modalities and fluorescent probes) (1). Agarose pads traditionally served this purpose (2–4), but in the last two decades have been largely superseded by microfluidic systems (5). A plethora of microfluidic designs have been utilized to generate a chemostat-like environment with continuous supply of fresh media and removal of excess cells (1, 6–20). Of these, the design that has proved the most popular involves polydimethylsiloxane (PDMS) “dead-end” growth channels oriented perpendicular to a main channel through which growth media is passed (9). It has earned the moniker “Mother Machine”, as a single “mother” cell at the end of each growth channel remains trapped throughout the experiment while all offspring eventually move into the main channel and exit the device (21). Originally used to study *Escherichia coli*, variations in the channel dimensions were quickly explored to accommodate the similarly rod-shaped *Bacillus subtilis* (22). Over a decade later, the Mother Machine remains the design of choice for most single-cell bacterial physiology studies. It has been used to study phenomena as diverse as single-cell aging (9, 23) and cell size homeostasis (24, 25) in steady-state conditions; gene expression (26), physiological growth (27), starvation adaptation (28) in time-varying environments; and antibiotic susceptibility (29), accumulation (30), and persistence (31). The ability to observe large numbers of individual cells in a controlled growth environment has enabled fundamental studies of phenotypic heterogeneity as a function of cell strain (32), growth condition (33), cell lineage (34), and cell age (35), as well as between species (*E. coli* and *B. subtilis*) (36).

Despite its numerous advantages, the Mother Machine has design constraints whose effects on qualitative and quantitative cell physiology merit closer examination. The loading procedure involves centrifugation of bacteria to force them into the dead-end channels, which may introduce stresses (37). Most studies have used a dedicated fluorescent reporter to identify and segment cells (26), but over-expression of a fluorescent marker can introduce a growth burden (38) and prolonged exposure to requisite bright illumination can lead to phototoxicity (39). Phase contrast microscopy provides a suitable alternative (40); although used less frequently due to challenges in detecting cells within the close walls of the Mother Machine, a number of tools have been developed for automated analysis of such data (41). Nonetheless, an extensive comparison of these tools noted systematic discrepancies in values obtained by different methods, necessitating care in interpretation of absolute quantities (42). In addition, unintended gradients in nutrient concentrations may well be unavoidable across each channel, and mechanical confinement in narrow growth channels may disrupt growth and morphology (43). That being said, growth of *E. coli* in suspension and in the Mother Machine have been found to be equivalent, on average (43). However, to fully characterize the effects of confinement on interpretation of quantitative Mother Machine studies, it is necessary to benchmark against an alternate single-cell technology that does not suffer from the same limitations.

Such a principled analysis has yet to be conducted, likely due to the lack of a suitable control that would permit the study of non-interacting, statistically identical, unconfined individual cells over multiple generations under constant environmental conditions. Unlike the aforementioned approaches, the SChemostat (14) design obviates the need for undesired confinement of cells to create a stable population for longterm imaging, instead relying on controllable surface adhesion to permit observations of a defined population of cells exposed to an identical flow environment within a relatively spacious microfluidic channel. Cell density may be tuned according to experimental considerations but is in general chosen to eliminate neighbor–neighbor contacts and maintain isolated microenvironments. This additionally assures the ability to acquire high-precision, high-resolution phase contrast images, which has enabled the study of cell cycle-dependent changes in cell morphology in the crescent-shaped *Caulobacter crescentus* (44). The technology has proved particularly fruitful for studying stochastic phenomena including scaling laws governing stochastic growth and division of individual cells (14, 45), stochastic intergenerational cell size homeostasis (46–48) and physiological adaption of single cells to precise time-dependent changes in growth conditions (49, 50). Because the SChemostat design eschews the need for overexpression of fluorescent reporters, cell centrifugation, surfactant treatment, or confinement that could result in mechanical stresses or heterogeneous microenvironments, it represents an ideal comparator to the Mother Machine for the measurement of baseline physiological parameters. We therefore developed experimental protocols to collect data from both the Mother Machine and SChemostat, under as similar conditions as possible, and quantified differences in single-cell growth and division dynamics between these approaches. Furthermore, we use these advances to address how the insights gleaned on cell size homeostasis of individual bacterial cells (46–48) are affected based on experimental and device modalities.

## Results

### Experimental Design

Wild-type *C. crescentus* cells exhibit a characteristic crescent shape, making them unsuitable for growth in the restricted environment of the Mother Machine, which is intended for rod-shaped cells. Deletion of crescentin, the intermediate filament-like protein responsible for imparting cell curvature, results in rod-shaped cells whose average growth rate is indiscernible from that of wild-type cells (51). Therefore, we generated a version of the *C. crescentus* strain used in SChemostat experiments (14) that lacks functional crescentin (ΔcreS), resulting in controllably adherent, rod-shaped cells suitable for growth in Mother Machine channels. In contrast to published protocols (9, 22), ultracentrifugation did not result in a significant improvement in cell loading efficiency; we therefore excised this step from the protocol in the interest of minimizing insults to cells prior to the experiment. We did however find it necessary to use the surfactant Pluronic F108, which reduces nonspecific surface interactions between PDMS channel walls and bacterial cells (52–54). Pretreatment of the device and supplementation of the growth media with Pluronic F108 (28, 55, 56) significantly reduced undesired adhesion to the PDMS, permitting observation of single cells over multiple generations without clogging of the growth channels.

Although many Mother Machine experiments are conducted using magnifications of 40× to 60× to maximize the amount of data acquired, this may result in reduced precision in the determination of cell sizes, which are required to accurately assess single cell growth rates. Therefore, we used the same imaging conditions standard for the SChemostat technology, consisting of a high-NA 100× objective (plus a 2.5× expander) to acquire high-resolution single-cell images. As *C. crescentus* cells divide more slowly than *E. coli* (approximately 70 min versus 20 min, respectively), images acquired at a frame rate of 1 min provide more-than-sufficient data points to precisely determine growth rates. To enable sideby-side comparison of data from the SChemostat and the Mother Machine, we modified the standard SChemostat image analysis routine to extract cross-sectional cell sizes from high-resolution, phase contrast images of cells growing in the Mother Machine. In conclusion, we established a protocol to successfully grow *C. crescentus* cells in the Mother Machine, and a routine to measure growth and division from phase contrast images (Fig. 1).

**Fig. 1.**
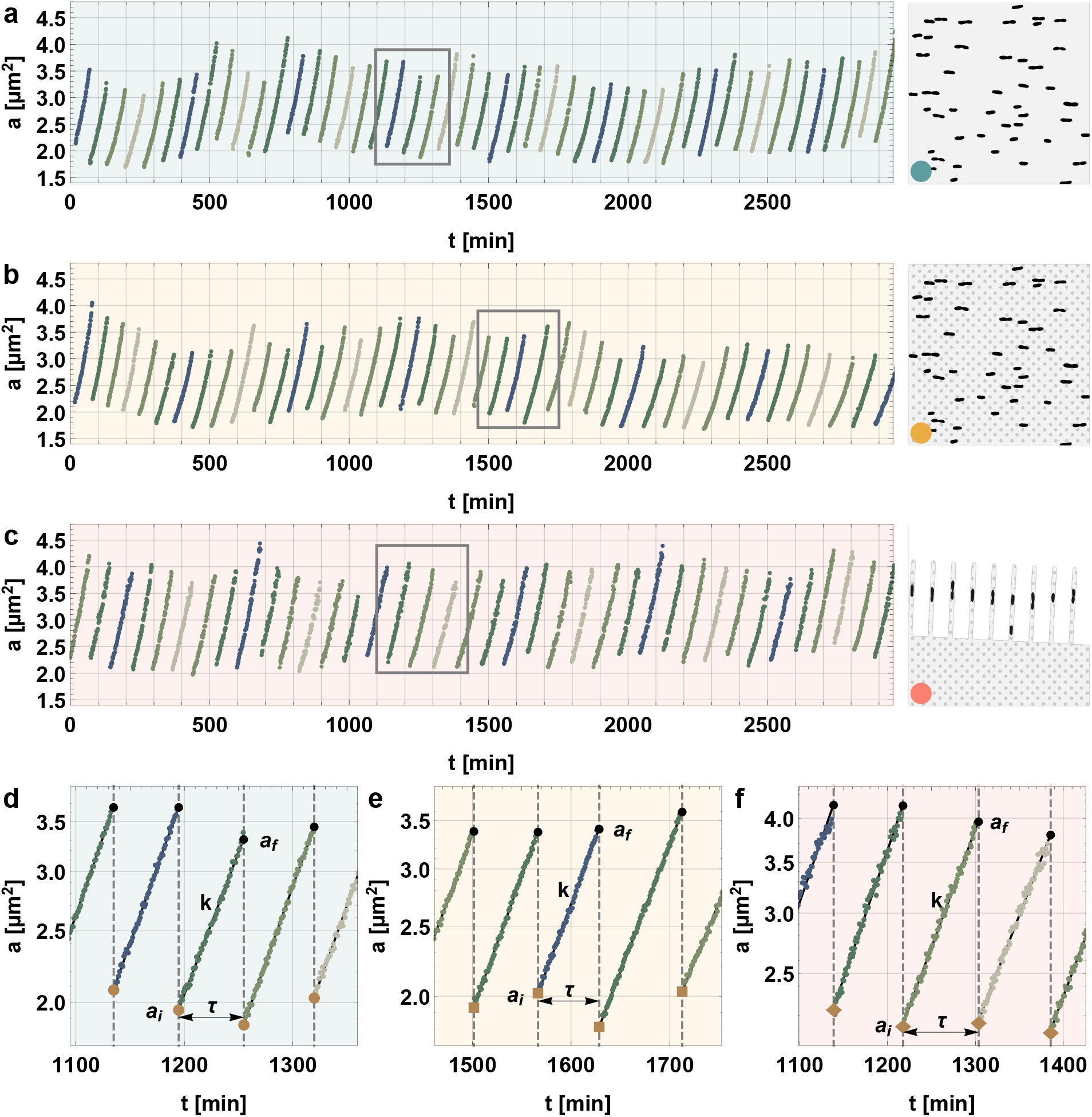
Representative Mother Machine and SChemostat data obtained using the reengineered modes of operation. **(a–c)** Time series of cell area over time for representative single *C. crescentus* cells from **(a)** SC^*\**^, **(b)** 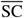, and **(c)** 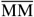. Insets at right show schematic representations of each condition as detailed in Fig. 4. **(d–f)** Zoomed insets of the time periods highlighted in **a–c**, respectively, on log-linear plots showing the definition of key growth and division parameters: initial area (*a*_*i*_), final area (*a*_*f*_), growth rate (*k*, given by the slope of the linear fit in the log-linear scale), and interdivision time (τ). SC^*\**^ : cells in SChemostat growing in complex media, 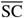: cells in SChemostat growing in complex media supplemented with Pluronic F108, 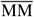: cells in Mother Machine growing in complex media supplemented with Pluronic F108.

Standardized growth and preparation of cells is necessary to compare experimental conditions between the SChemostat and Mother Machine platforms (see Fig. 2a). A key difference in the established protocol involves Pluronic F108 pretreatment of the Mother Machine prior to, and media supplementation during, experiments. However, the effects of Pluronic F108 on *C. crescentus* growth are unknown. To control for Pluronic F108, we observed ΔcreS cells growing in the SChemostat in complex media (SC^*\**^), cells growing in the SChemostat in complex media supplemented with Pluronic F108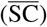, and cells growing in the Mother Machine in complex media supplemented with Pluronic F108 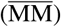. Finally, to minimize potentially confounding effects due to nutrient depletion or mechanical pressure resulting from neighboring cells within the Mother Machine, here we use data obtained from growth channels occupied by a single *C. crescentus* cell (Fig. 2b–d).

**Fig. 2.**
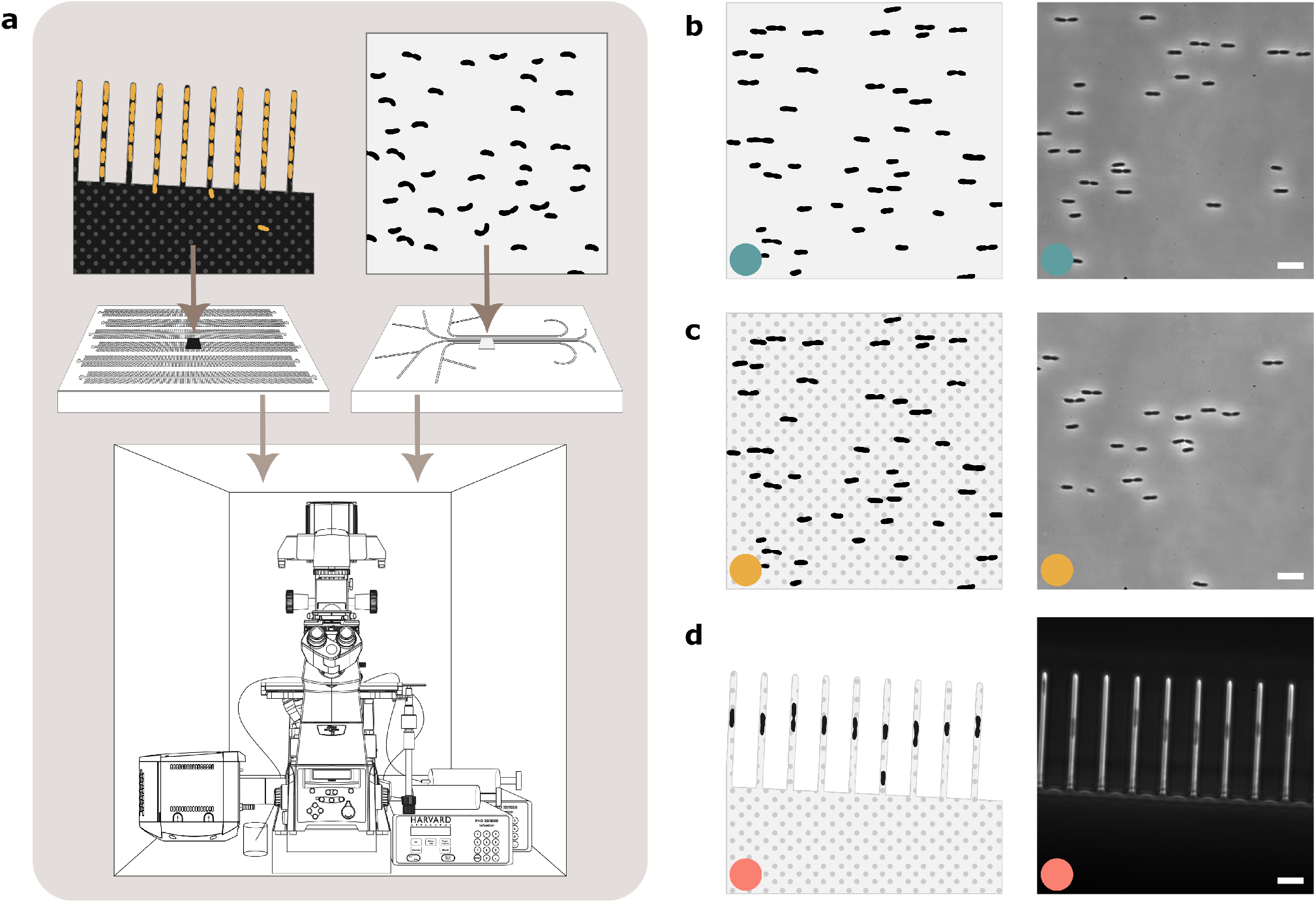
Comparison of Mother Machine and SChemostat experiment protocols in both standard and reengineered modes of operation. **(a)** Schematic of typical experiments conducted in the Mother Machine (*left*) and SChemostat (*right*) setups. Standard Mother Machine data are collected from rod-shaped cells using fluorescence microscopy in the presence of a molecule to reduce sticking (BSA or Pluronic F108); standard SChemostat data are collected from any-shaped cells using phase contrast microscopy after transiently induced surface adhesion. **(b–c)** Overview of experimental conditions used in this study, showing schematic representation (*left*) and representative images (*right*) for **(b)** SC^*\**^, **(c)** 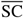, and **(d)** 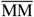 cells. Scale bars: 5 μm. SC^*\**^ : cells in SChemostat growing in complex media, 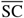: cells in SChemostat growing in complex media supplemented with Pluronic F108, 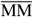: cells in Mother Machine growing in complex media supplemented with Pluronic F108.

**Fig. 3.**
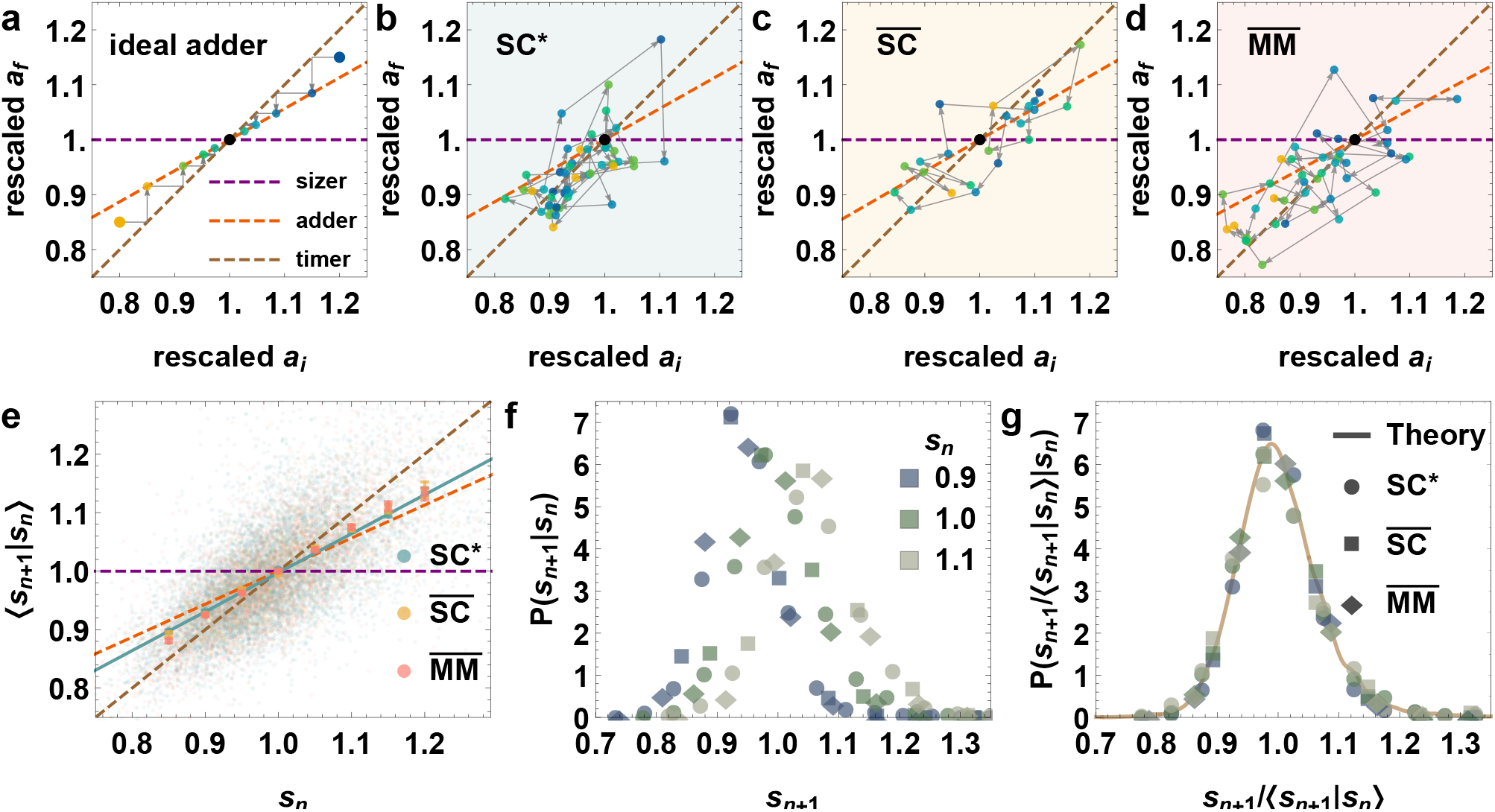
The inherently stochastic nature of inter-generational size dynamics. **(a–d)** Typical intergenerational final size (*a*_*f*_) versus initial size (*a*_*i*_) trajectories are plotted for **(a)** two idealized “theoretical” cells following the adder model, **(b)** an experimentally observed cell in SC^*\**^, **(c)** an experimentally observed cell in 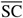, and **(d)** an experimentally observed cell in 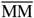. The initial and final sizes are rescaled by their respective population mean values. The traditional quasi-deterministic adder–sizer–timer paradigm of homeostasis in **(a)**, marked as dashed lines, evidently proves inadequate in capturing the direct experiment measurements in **(b–d)**, necessitating the fully stochastic description (Eq. (3)) consistent with observations in **(e–g). (e)** Binned mean of the next generation’s mean-rescaled initial size (*s*_*n*+1_) plotted as a function of the current generation’s mean-rescaled initial size (*s*_*n*_) for SC^*\**^ (teal), 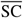 (yellow), and 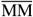 (pink) cells (bars indicate standard deviations). The background scatter shows individual pairs of experimentally obtained *s*_*n*_,*s*_*n*+1_ values. The teal line is the linear fit to the SC^*\**^ data, which also fits the other two conditions well. **(f)** Experimentally measured conditional distributions of the next generation’s mean-rescaled initial size (*s*_*n*+1_) given the current generation’s mean-rescaled initial size (*s*_*n*_), plotted for three different values of *s*_*n*_ for SC^*\**^ (circle), 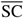 (square), and 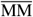 (diamond) cells. **(g)** The distributions in (f), when rescaled by their respective mean values, become independent of *s*_*n*_ and growth condition. The theoretical curve is fit to SC^*\**^ data, but it matches all three conditions well. SC^*\**^ : cells in SChemostat growing in complex media, 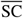: cells in SChemostat growing in complex media supplemented with Pluronic F108, 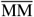: cells in Mother Machine growing in complex media supplemented with Pluronic F108.

The size growth of a single *C. crescentus* cell under balanced growth conditions over one generation is exponential, 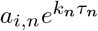, where *a*_*i*_ and *a*_*f*_ are the initial and final cell areas, respectively, *k* is the growth rate, *·* is the interdivision time and the subscript indicates the generation number (14). We also calculate the division ratio *r*_*n*_ = *a*_*i,n*+1_*/a*_*f,n*_, which gives the fraction of the total area of the mother stalked cell immediately before division inherited by the daughter stalked cell immediately after division. We find that the asymmetric division of *C. crescentus* cells ensures that irrespective of the initial alignment of the cell trapped at the end of the Mother Machine growth channel, starting from the second generation onwards, the eponymous mother cell is always stalked, with the stalk oriented toward the closed end of the channel. Since imaging and data acquisition start a sufficient time after device loading, all data presented here are thus collected from a defined population of stalked cells within both the Mother Machine and SChemostat approaches.

### Glossary

For convenience in interpreting the results presented here, we include the following glossary to distinguish between the three distinct device and experimental configurations considered here:

- **SC***\** : ΔcreS cells growing in the SChemostat in complex media
- 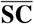 : ΔcreS cells growing in the SChemostat in complex media supplemented with surfactant (Pluronic F108)
- 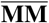 : ΔcreS cells growing in the Mother Machine in complex media supplemented with surfactant (Pluronic F108)

### The interdivision time distribution has larger mean and variability for cells under mechanical confinement in the Mother Machine setup

Interestingly, while the population-averaged single-cell growth rates of 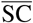 and 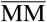 cells are reduced by ∼12% compared to SC^*\**^ cells, the 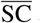 and 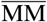 single-cell growth rate distributions are remarkably similar. We deduce that the addition of the surfactant (Pluronic F108), a necessary step for the Mother Machine protocol, results in the observed differences in growth rate distributions (Fig. 4a).

**Fig. 4.**
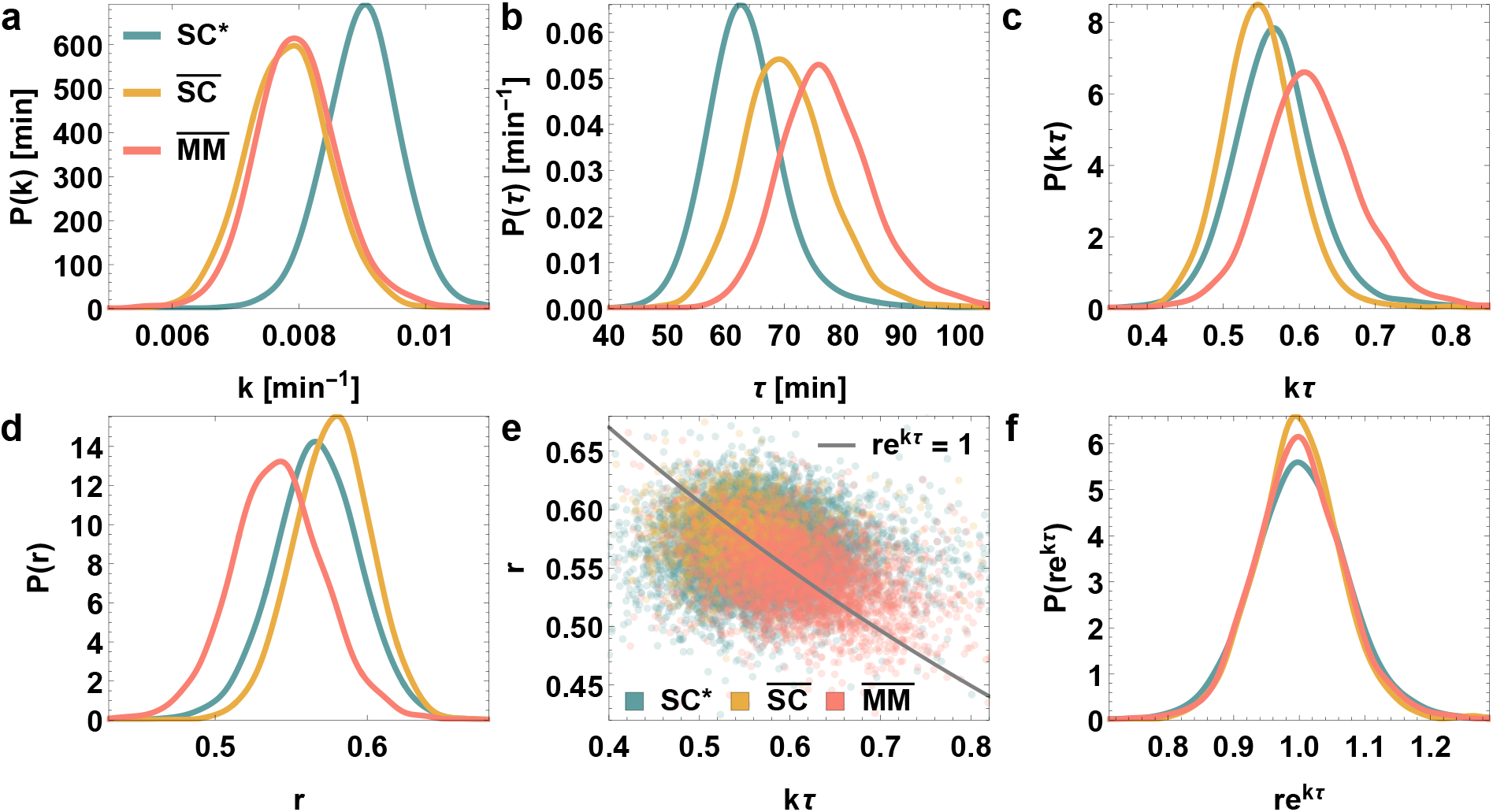
Cells adjust division ratio to balance changes in growth rate and interdivision time. Distributions of growth and division variables are plotted for SC^*\**^ (teal), 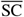 (yellow), and 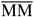 (pink) conditions. **(a)** Growth rate (*k*) distributions are identical betwen 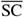 and 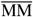, and both lower on average than SC^*\**^ . **(b)** Interdivision time (*τ*) distributions differ betwen each condition, with 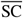 slower than SC^*\**^, and 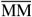even slower. **(c)** Distributions of *kτ·* and **(d)** division ratio (*r*) show opposite trends, such that **(e)** the values of *r* and *kτ·* match *re*^*kτ·*^ = 1 and **(f)** *re*^*kτ·*^ distributions are roughly the same irrespective of the growth condition. Thus, division ratio (*r*) compensates for differences in *kτ·* . SC^*\**^: cells in SChemostat growing in complex media, 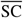: cells in SChemostat growing in complex media supplemented with Pluronic F108, 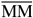: cells in Mother Machine growing in complex media supplemented with Pluronic F108.

Intriguingly, for cells in the SChemostat, the reduction in growth rate due to Pluronic F108 is compensated by an ∼11% increase in interdivision time (Fig. 4b), such that the distributions of their product, *kτ*, for SC^*\**^ and 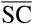 are very similar (Fig. 4c). In contrast, 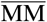 cells exhibited a greater increase in interdivision time than expected due to the surfactant (Pluronic F108) alone (∼22% increase compared to SC^*\**^ and ∼10% compared to 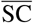), indicating that additional factors slow down the time to cell division in the Mother Machine (Fig. 4b). This additional delay in the division process ensures that the *k·τ* distribution of 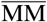 cells is distinctly shifted toward a larger mean value and has a greater spread (variance), indicating increased variability compared to both SC^*\**^ and 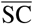 cells (Fig. 4c). The coefficients of variation (CVs) of these quantities are reported in Supplementary Table 2.

To further investigate whether this additional increase in interdivision time within the Mother Machine could result from reduced nutrient diffusion within the growth channels, we compared the distributions of relevant growth and division variables for cells growing in different channel lengths in the Mother Machine. We observed no significant differences between these distributions across channel lengths, implying that mechanical confinement, and not nutrient diffusion as a function of channel length, causes the additional delay in cell division in our Mother Machine setup (with one cell per channel).

### Variation in *kτ* across device and experimental configurations is compensated by complementary variation in the division ratio

Despite differences in the distributions of *k, τ* and *r* (Fig. 4a–d), the distributions of *re*^*kτ*^ (where *r* is the division ratio) are remarkably similar for all three variations of the experimental setup (Fig. 4e–f). Furthermore, the coefficient of variation (CV) of *re*^*kτ*^ is less than the CV of either *k* or *τ·* (see Supplementary Table 2), indicating that fluctuations in *r* and *kτ* are anti-correlated. The value of *re*^*kτ*^ corresponding to a given generation is equal to the the ratio of the next generation’s initial size (size at birth) to the current generation’s initial size, since

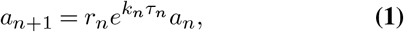

where the subscripts denote generation numbers and *a* denotes the initial size. The distribution of *re*^*kτ*^ must be approximately balanced around 1 to maintain cell size homeostasis (i.e., to prevent runaway cell sizes). Otherwise, if these values are consistently greater (smaller) than 1 across generations, initial sizes will increase (decrease) indefinitely, breaking intergenerational cell size homeostasis. Accordingly, we observe that the population-wide distributions of *re*^*kτ*^ are centered around 1. A similar finding has been reported for *E. coli* cells in a study that looked at individual lineages (57).

### Emergent simplicities governing stochastic intergenerational homeostasis of mean-rescaled cell sizes are universal across device and experimental configurations

The straightforward requirement for homeostasis that distributions of *re*^*kτ*^ must be centered around 1 cannot explain why the entire *re*^*kτ*^ distributions are observed to match across growth conditions (Fig 4f). Instead, we find that this observation is a consequence of a remarkable emergent simplicity: cells in all three variations of the experimental setup follow *the same* precision kinematics of stochastic intergenerational cell size homeostasis.

Given a generation’s initial cell size, the next generation’s initial size can be obtained by multiplying by the current generation’s value of *re*^*kτ*^ (see Eq. (1)). Thus, if the intergenerational scaling factor, *re*^*k·τ*^, follows the same intergenerational dynamics across different devices and conditions (consistent with Fig. 4f), we can expect the intergenerational dynamics of initial sizes to also be identical. This clearly does not hold for 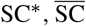, and 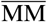 cells (Fig. 5). Instead, when the steady state initial size distributions are rescaled by their respective mean values, we find the emergent simplicity that the resulting distributions are identical for 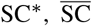, and 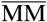 cells (Fig. 5). We bridge this discrepancy through the following proposition. We know that *re*^*kτ*^ in a given generation depends on that generation’s initial size (46), otherwise it would result in ‘timer’-like behavior, which breaks cell size homeostasis (47, 58, 59). Based on the observations that 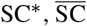, and 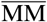 cells have the same *re*^*kτ*^ distributions but different steady state initial size distributions and same mean rescaled steady state initial size distributions, we propose that *re*^*kτ*^ must depend on the *rescaled* initial size instead, rescaled by a scaling factor that is proportional to the population-averaged initial size for the particular growth condition. This allows for the absolute cell sizes to vary across growth conditions (through the difference of a constant scaling factor) while following identical laws determining cell sizes over successive generations. Indeed, despite cells in 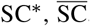, and 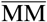 conditions having different distributions of initial and final sizes (Fig. 5a,c), when the steady-state size distributions are rescaled by their respective mean values, we find that the resulting distributions are identical for 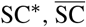, and 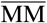 cells (Fig. 5b,d)! This remarkable empirical observation implies that once appropriately rescaled, cell sizes in all three conditions have the same homeostatic distributions.

**Fig. 5.**
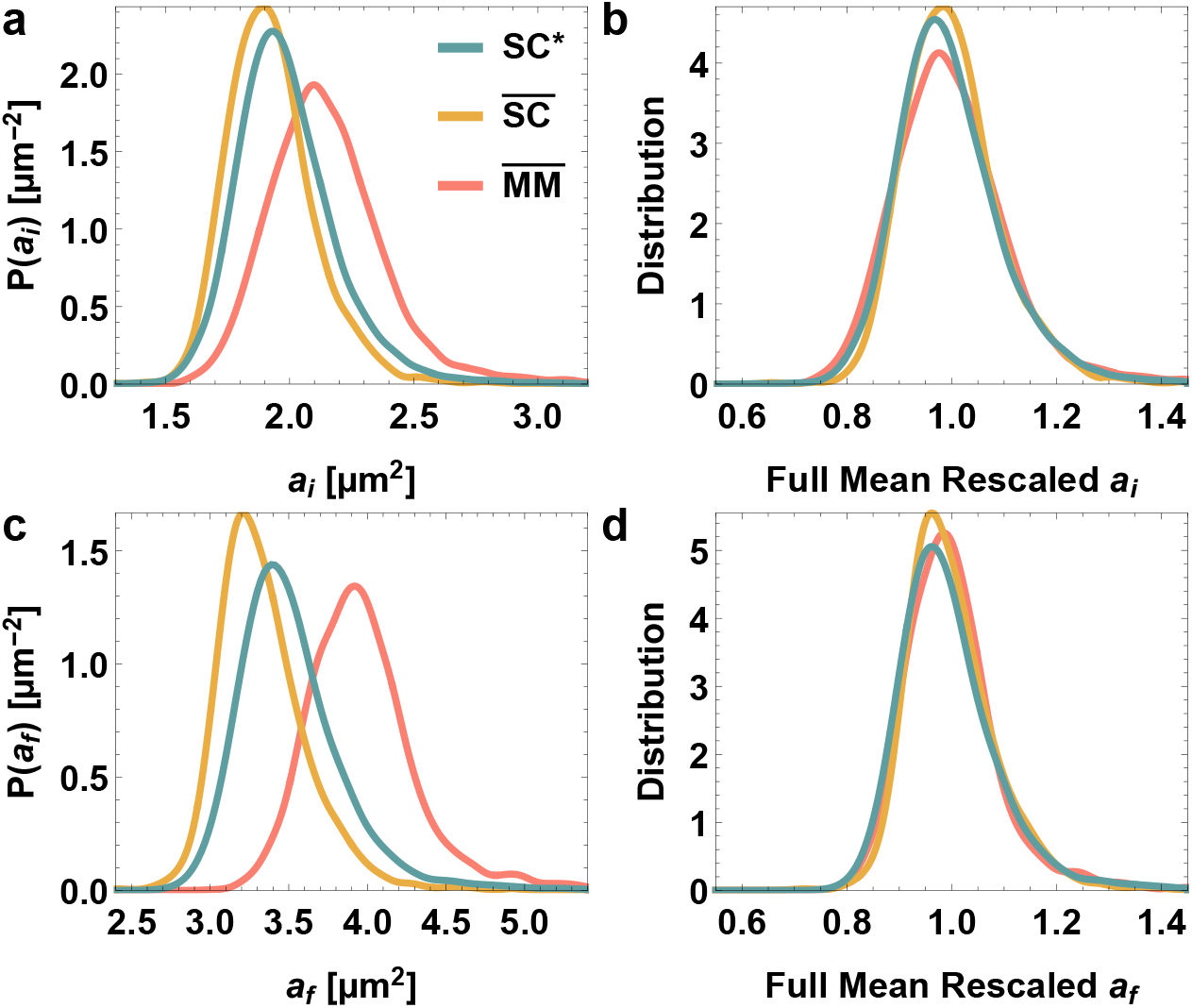
Steady state cell size distributions at birth (initial areas) and division (final areas) along with their mean-rescaled counterparts in different experimental platforms and growth conditions. **(a)** Steady state distributions of initial area (area at birth) are shown for SC^*\**^ (teal), 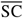 (yellow), and 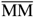 (pink) cells. **(b)** The distributions in **a** rescaled by their respective mean values are remarkably similar across platforms and growth conditions. **(c)** Steady state distributions of final area (area at division) are shown for 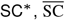, and 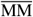 cells. **(d)** The distributions in **c** rescaled by their respective mean values are also remarkably similar across experimental platforms and growth conditions. SC^*\**^ : cells in SChemostat growing in complex media, 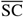: cells in SChemostat growing in complex media supplemented with Pluronic F108, 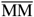: cells in Mother Machine growing in complex media supplemented with Pluronic F108.

We track the intergenerational evolution of initial cell sizes rescaled by the population mean of the corresponding experiment denoting the rescaled cell size by *s*. We now turn to the emergent simplicities in the stochastic intergenerational dynamics of cell sizes. Traditionally, deterministic models corresponding to the idealizations of either sizer (constant final size), adder (constant difference between final and initial sizes), or timer (constant ratio of final to initial sizes) have been used to characterize cell size homeostasis in a range of unicellular organisms, notably the adder model for *E. coli* and *B. subtilis* (24, 58, 60–62). However, the behaviors of the mythical average cell prove inadequate in capturing basic phenomenologies of stochastic intergenerational cell size homeostasis, including the *inter*-generational scaling law we previously reported in (46–48) (see Fig. 3).

*C. crescentus* in particular deviates substantially from the adder-sizer-timer framework (46, 61) but the fully stochastic behavior is precisely captured by the theoretical framework we have presented in (47) (Fig. 3a-d). This theory accurately predicts the distribution of initial sizes after *n* generations for a cell having initial size *a*_0_ in the starting (zeroth) generation (46). Accordingly, this distribution is given by:

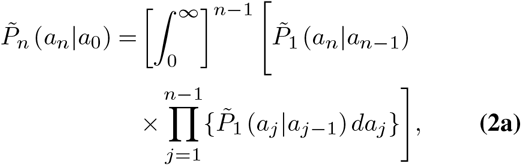

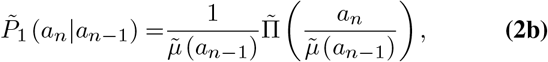

where 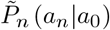 is the growth condition-dependent conditional distribution of possible initial sizes after *n* generations, given that the initial size in the zeroth generation is *a*_0_. 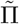 is the mean-rescaled distribution of the next generation’s initial size given the current generation’s initial size. That it is independent of the current generation’s initial size is the relevant emergent simplicity (46–48). 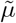is the conditional average of the next generation’s initial size given the current generation’s initial size; it is observed to be a linear function of the current generation’s initial size (46, 48). Rewriting Eq. (2) in terms of the mean rescaled initial size, *s* = *a/ā* with *ā* being the population mean of *a*, the conditional distribution after *n* generations becomes:

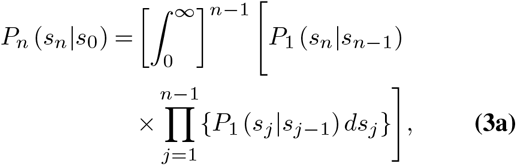

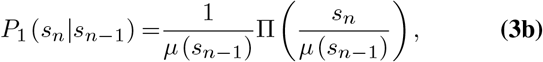

where,

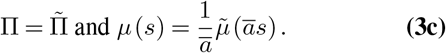

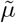 and thus 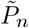 are different for different growth conditions. Remarkably then, once we mean-rescale by the population means, we find that all growth conditions have the same values of *μ* and r (Fig. 3e–g), and hence the same *P*_*n*_. We establish this by comparing the experimentally obtained conditional distribution of mean-rescaled initial sizes after *n* generations, given the mean-rescaled initial size in the zeroth generation, for 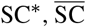, and 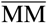 cells, and using the theoretical predictions for *P*_*n*_ for the SC^*\**^ cells given by Eq. (3). (We note that 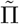and 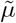 are experimentally measured; there are no fine-tuning fitting parameters). The observed and calculated distributions match compellingly for different values of final generation, *n*, and initial rescaled size, *s*_0_ (Fig. 6). In particular, they converge to the same homeostatic distribution after sufficient generations have elapsed.

**Fig. 6.**
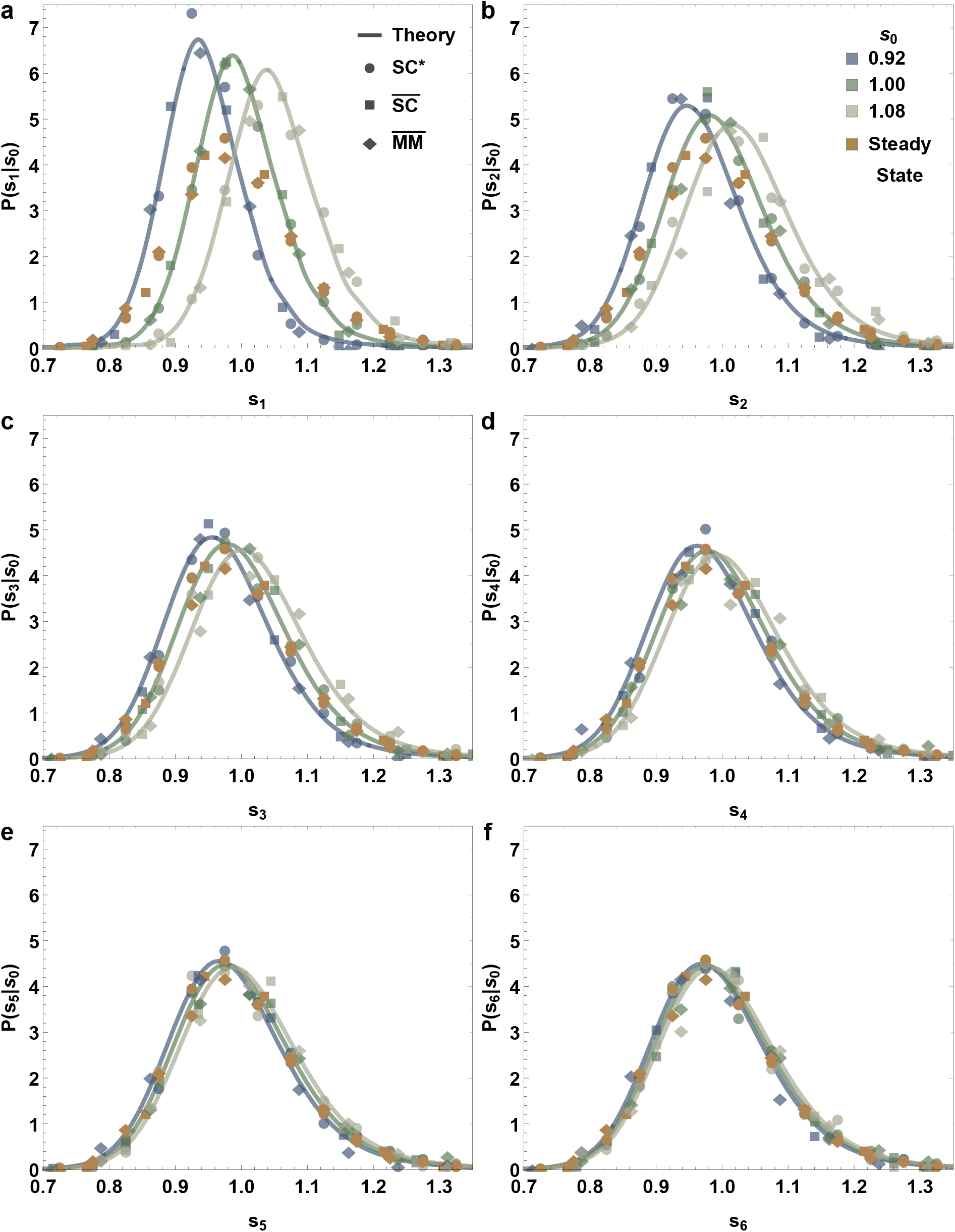
Emergent simplicities in the stochastic intergenerational homeostasis of mean-rescaled sizes-at-birth (initial cell areas) across experimental platforms and conditions. For cells starting from different given mean-rescaled initial areas (different values of *s*_0_, marked by different colors), the distributions of mean-rescaled initial areas after successive generations (*s*_*n*_) are shown, for *n* going from 1–6 in (**a–f**), respectively. The distributions for different experimental conditions are plotted with different markers for SC^*\**^ (circle), 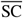 (square) and 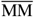 (diamond). The solid lines are the theoretical predictions for the SC^*\**^ data, but nevertheless match all three experimental conditions. The distributions in gray in each panel are the experimentally obtained steady state mean-rescaled initial area distributions. All distributions converge to these steady state distributions as *n* increases, irrespective of the starting *s*_0_, indicating that cell size is in homeostasis. SC^*\**^ : cells in SChemostat growing in complex media, 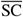: cells in SChemostat growing in complex media supplemented with Pluronic F108, 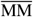: cells in Mother Machine growing in complex media supplemented with Pluronic F108.

Thus, we conclude that despite apparent differences in the distributions of interdivision times, growth rates, division ratios and cell sizes, once cell sizes are appropriately rescaled by the corresponding population averages, they undergo identical stochastic intergenerational dynamics for cells growing in all three modalities: 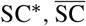, and 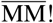 The significant implication is that the emergent simplicities previously encountered in the context of intergenerational homeostasis of cell sizes (46–48) are conserved across these devices and conditions.

## Discussion

The results presented here provide insights into the nature of the robustness of the bacterial growth and division machinery. Despite significant changes in individual growth and division parameters, the determinant for cell size homeostasis, *re*^*kτ*^, remains invariant. Furthermore, the fact that the distributions of *re*^*kτ*^ across devices and conditions themselves align so closely implies that the emergent simplicities governing stochastic intergenerational homeostasis of mean-rescaled cell sizes are universal across device and experimental configurations! With knowledge of this remarkable new emergent simplicity, principled quantitative comparisons of stochastic growth and division dynamics measured in different microfluidic devices is made possible. Followup studies in different growth media or at different temperatures, pH, or other growth conditions could serve to further broaden the scope of the empirical findings reported here.

Mechanical confinement is known to affect growth; channels that are too narrow lead to anomalous morphology and larger-than-expected volumes in *E. coli* (63). Even in standard size channels, it has been observed that *E. coli* adapt to the confined environment, approximately preserving cell volume but adjusting aspect ratio by becoming narrower and longer than cells grown in suspension (43). Mechanical forces opposing cell growth (such as friction due to channel walls) can decrease elongation rate in *E. coli* (64) and may constitute the main factor limiting cell growth in long narrow channels, even though growth rates between standard Mother Machine conditions and bulk culture are, on average, equivalent (43). We attribute reduced growth rate of *C. crescentus* in the Mother Machine to the presence of Pluronic F108; however, this does not account for deviations in the interdivision time distribution. These results demonstrate the need to fully characterize the effects of environmental constraints on observed phenomena, especially when moving from population-level aggregate to stochastic singlecell measurements and attempting to draw inferences from individual-level variations. They also serve to underscore that the mythical ‘average cell’ does not realistically capture the complex behaviors and stochastic dynamics of the individual cell, as is increasingly appreciated in different biological contexts (65–72). Direct genetic perturbations to division machinery such as FtsZ (25) and Min (73) have been demonstrated to alter division timing independent of growth rate, but future studies examining the dynamics of specific components of the division machinery are needed to characterize the relationship between mechanical confinement and disruptions in the timing of cell division.

Scaling laws are ubiquitous in nature, and have been reported in the context of fluctuations in growth and division dynamics of different bacterial species. These scaling laws offer insights into the mechanisms underlying growth and division. Previously, *intra*generational scaling laws have been reported for many microorganisms (74). In particular, the mean-rescaled cell interdivision time distribution (from different growth conditions) (14), the mean-rescaled cell size distributions (from different times since the last division event) (14), and the mean-rescaled cell age distributions (from different growth conditions) (45, 50) undergo scaling collapses for *C. crescentus* cells in balanced growth conditions. Other known examples of *intra*generational scaling laws include the mean-rescaled distributions of initial size, final size, growth rate, and division time across growth conditions for *E. coli* cells growing in the Mother Machine (24). In contrast, here we focus on *inter*generational scaling laws, generalizing further the ongoing line of work reported in (46–48) by drawing comparisons across *different experimental setups* and identifying invariant behaviors. In particular, we find that the mean-rescaled *intra*generational growth rate and division time distributions are not invariant across devices but, remarkably, the initial and final cell size distributions still are. This *inter*condition scaling law, when paired with the invariance of *re*^*kτ*^ across devices, results in the general validity across experimental realizations of the *inter*generational scaling law governing cell size homeostasis (Fig. 3).

## Materials and Methods

### Experimental Setup

SChemostat and Mother Machine microfluidic devices were fabricated from polydimethylsiloxane (PDMS) using standard soft lithography techniques. Mother Machine devices were incubated with 0.1 g/mL Pluronic F108 (Sigma-Aldrich, 542342) for 24 h prior to each experiment. *C. crescentus* cells were grown to log phase at 30°C in peptone yeast extract (PYE) liquid cultures. For SC^*\**^ and 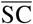 experiments, cells were exposed to 75 μM vanillic acid (Sigma Aldrich, H36001) for 75 min immediately prior to loading into the device. For all experiments, cells were loaded into the device and incubated for 60 min either in PYE (SC^*\**^) or in PYE supplemented with 1% of 0.1 g/mL Pluronic F108 (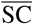 and 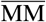). The same growth media was then perfused through the device at 10 μL/min and phase contrast images were captured every 60 s, as described previously (46).

### Image Analysis

Chemostat data were analyzed as previously described (46). To adapt this routine to phase contrast images obtained in the Mother Machine, we generated a background image for each movie from a series of registered images sampled over time, then subtracted this background from each image to remove features corresponding to the device. This procedure provided images suitable for pixel-based segmentation as previously described (46).

## Acknowledgments

We thank Purdue University Startup funds, Purdue Research Foundation, the Purdue College of Science Dean’s Special Fund, and the Showalter Trust for financial support. K.F.Z., K.J., and S.I.-B. acknowledge support from the Ross-Lynn Fellowship award. K.J. and S.I.-B. acknowledge support from the Bilsland Dissertation Fellowship award. W.C. and S.I.-B. thank the Louis Stokes Alliances for Minority Participation (LSAMP) for financial support. S.R. and S.I.-B. acknowledge financial support from the Data Mine Learning Community at Purdue. S.I.-B. thanks the Harvard Medical School’s Department of Systems Biology and Johan Paulsson for graciously hosting her as an extended visitor during early stages of this work. K.F.Z. and S.I.-B. thank members of the Johan Paulsson lab, in particular Silvia Canas Duarte, Brandon Seo, and Carlos Sanchez, for invaluable insights on Mother Machine device design and fabrication.

## Contributions

S.I.-B. conceived of the research; K.F.Z., K.J., C.S.W., R.R.B., and S.I.-B. designed the research; K.F.Z. spearheaded engineering and technology development under the guidance of S.I.-B.; K.J. performed data analysis and theory under the guidance of S.I.-B.; K.J., R.R.B., and S.I.-B. performed and checked analytic calculations; K.F.Z. and S.I.-B. performed experiments; C.S.W. spearheaded image analysis efforts and wrote the automated analysis codes for both setups presented here; S.R. contributed to a preliminary version of image analysis of Mother Machine data with input from K.J. and C.S.W.; S.R. manually supervised image analyses with inputs from K.J.; W.C. contributed to manual supervision of image analyses; K.F.Z., K.J., C.S.W., R.R.B., and S.I.-B. wrote the paper; S.I.-B. supervised the research.

## Notes

### Competing Interest Statement

The authors have declared no competing interest.

### Summary of Updates

The manuscript has been updated to improve presentation and clarity.

